# Healthy aging, processing speed, and mnemonic brain state engagement

**DOI:** 10.64898/2026.04.28.721318

**Authors:** Hannah R. Buras, Subin Han, Nicole M. Long

## Abstract

Healthy older adults exhibit both selective impairments in episodic memory – memory for events situated within a specific time and place – and deficits in executive function, reflected by difficulty switching between different tasks and inhibiting task-irrelevant information. Prior work has shown that older adults show diminished mnemonic brain state engagement – recruitment of whole brain activity patterns that selectively support memory encoding and memory retrieval. Our hypothesis is that older adults are biased toward the retrieval state and, due to executive function deficits, cannot easily switch out of this state when task-irrelevant. Our goal was to determine the extent to which stimulus processing time impacts older adult mnemonic state engagement, with the expectation that longer processing times would enable older adults to switch out of a task-irrelevant retrieval state. We recorded scalp electroencephalography (EEG) while younger and older adult participants explicitly encoded and retrieved object stimuli under variable stimulus durations. Using a combination of multivariate decoding approaches, we find that under time constraints, older adults both under-recruit a young-adult like retrieval state when task-relevant, but over-recruit a participant-specific retrieval state when task-irrelevant. Older adults may thus recruit idiosyncratic activity patterns to compensate for difficulties engaging young-adult like mnemonic brain states. Taken together, these findings suggest that although older adults retain the ability to engage encoding and retrieval brain states, they require more processing time to both initiate and maintain goal-relevant mnemonic states.

## 1 Introduction

> *That was how the old ended up not remembering recent events–by not seeing them in the first place. Memory lost before it ever came to be, because one was focusing so intently on the past.*
>
> — Kim Stanley Robinson, *Green Mars*

Healthy older adults typically demonstrate selective memory declines with impaired episodic memory – memory for events situated within a specific time and place – relative to younger adults (Tulving, 1983; Wingfield & Kahana, 2002; Golomb, Peelle, Addis, Kahana, & Wingfield, 2008). For example, an older adult may fail to remember a specific trip taken to the local library last Tuesday. Aging-related declines in episodic memory may be driven by an increased reliance on semantic memory – memory for general knowledge or facts (Tulving, 1972; Brod & Shing, 2019; Wynn, Ryan, & Moscovitch, 2020; Matijevic, Hoscheidt, & Ryan, 2024). This reliance on semantic memory may be connected with an age-related bias to engage pattern completion, the reactivation of stored memory traces based on partial cues (McClelland, McNaughton, & O’Reilly, 1995; Vieweg, Stangl, Howard, & Wolbers, 2015). We have previously found evidence to suggest that older adults over-recruit the retrieval state, a brain state that supports internal processing to access stored information, at the expense of engaging the encoding state, a brain state that supports the formation of new memories (Moore, Smith, & Long, 2025). Given that older adults have difficulty switching between different tasks and inhibiting task-irrelevant information, including inhibiting retrieval of semantic information (Hamm & Hasher, 1992; Kray & Lindenberger, 2000; Wynn et al., 2020), age-related changes in mnemonic state engagement may be due to alterations in executive processing (Reuter-Lorenz & Park, 2014). Such alterations may be revealed by varying the time allotted to engage task-relevant mnemonic states and inhibit task-irrelevant mnemonic states. The aim of the current study is to determine the extent to which stimulus processing time impacts older adult mnemonic state engagement.

Older adults’ episodic memory impairments may be the result of biases toward semantic knowledge. There is an overall shift toward semantic memory across the lifespan (Ofen & Shing, 2013) and older adults over-rely on prior knowledge (Wynn et al., 2020). The recall of episodic vs. semantic details are negatively correlated, an effect which increases with age (Devitt, Addis, & Schacter, 2017). Orienting tasks that focus attention specifically to semantic information can lead to decreased free recall performance in younger adults and particularly reduce reliance on episodic (temporal) information shared between study items (Long & Kahana, 2017). Relative to younger adults, older adults show less reliance on temporal associations during free recall tasks (Wingfield & Kahana, 2002; Golomb et al., 2008). Greater dependence on accumulated semantic knowledge may thus be compensatory for altered episodic mechanisms (Castel, 2005); however, dependence on semantic knowledge can become maladaptive when prior knowledge conflicts with current task demands (Lalla, Tarder-Stoll, Hasher, & Duncan, 2022).

Aging-related changes in cognition have been linked to alterations in the default mode network (DMN, Raichle et al., 2001). The DMN is a brain network that supports internally-directed mentation (Buckner & DiNicola, 2019), including semantic processing (Binder, Desai, Graves, & Conant, 2009) and accessing prior knowledge (Rubin, 2021). Both activity and connectivity within the DMN change with age and are associated with changes in episodic memory performance (Grady, Springer, Hongwanishkul, McIntosh, & Winocur, 2006; Damoiseaux, 2017; Staffaroni et al., 2018). Over-engagement of internal processing via the DMN can limit external processing of to-be-remembered stimuli leading to episodic memory deficits (Stevens, Hasher, Chiew, & Grady, 2008; Kim, 2011). According to the default-executive coupling hypothesis of aging (DECHA, Turner & Spreng, 2015; Spreng & Turner, 2019), the DMN becomes more persistently engaged and more strongly coupled with the executive control network in healthy aging. This shift may reflect a compensatory response to declining control resources, whereby prior knowledge is recruited to support task performance. Indeed, default-executive coupling has been shown to facilitate the process of integrating new information with existing representations, particularly when the new information is consistent with prior knowledge (Amer, Giovanello, Nichol, Hasher, & Grady, 2019). However, when prior knowledge is irrelevant to the current task, according to the DECHA, this inflexible coupling can hinder performance (Rieck, Rodrigue, Boylan, & Kennedy, 2017; Turner & Spreng, 2015). Failure to sufficiently suppress DMN activity can allow task-irrelevant thoughts and internal processes to intrude during encoding thus increasing distraction and leading to diminished memory performance (Amer, Anderson, Campbell, Hasher, & Grady, 2016; Sambataro et al., 2010). Given older adults’ propensity to over-rely on prior knowledge, coupled with hyperactivity of the DMN, older adults may have an increased likelihood of entering a task-irrelevant retrieval state which can worsen episodic memory formation.

Older adults also show declines in executive function – a set of interrelated cognitive processes including working memory, inhibitory control, and attention – relative to younger adults. Healthy aging has been tied to overall processing speed deficits (Salthouse, 1993; Park et al., 1996), an effect which often increases with task difficulty and switching between tasks (Salthouse, Fristoe, McGuthry, & Hambrick, 1998). In particular, older adults show greater task-switching costs compared to younger adults, with worse behavioral performance (lower accuracy, slower response times) when they must switch between two different tasks and ignore previously relevant, now irrelevant, information (Kray & Lindenberger, 2000). Older adults display impaired control over inhibitory processes which can result in retrieval and maintenance of task- irrelevant knowledge (Amer, Wynn, & Hasher, 2022; Lustig, Hasher, & Zacks, 2007), with a particular bias for semantic information (Wynn et al., 2020). Older adults over-maintain previously activated, but now irrelevant, information, creating “cluttered” memory representations, increased susceptibility to interference, and competition at retrieval (Biss, Ngo, Hasher, Campbell, & Rowe, 2013; Spreng & Turner, 2019). This reduced suppression is also associated with persistent, incidental processing of distractors thereby contributing to the “clutter” and worsening memory performance (Amer & Hasher, 2014). Consistent with these behavioral findings, neuroimaging evidence suggests that age-related declines in executive processes are associated with alterations in large-scale brain networks that support cognitive control (Rieck, Baracchini, & Grady, 2021). In particular, older adults show reduced modulation of frontoparietal control regions and diminished suppression of the DMN as task demands increase, as well as stronger coupling between lateral prefrontal cortex and the DMN during challenging control tasks (Grady et al., 2006; Stevens et al., 2008). These patterns suggest that age-related changes in executive control may be linked with alterations in mnemonic brain state engagement, which together impact memory performance.

Mnemonic brain states support remembering in younger adults. Memory encoding and memory retrieval constitute neurally distinct brain states – whole brain activity/connectivity patterns (Harris & Thiele, 2011) – that are engaged in response to demands to form new memories and access stored memories, respectively (Long & Kuhl, 2019; Smith, Moore, & Long, 2022). Encoding and retrieval rely on overlapping, but opposing neural circuit mechanisms (Hasselmo, Bodelón, & Wyble, 2002; Hasselmo, 2005) such that the two mnemonic states tradeoff. Using a mnemonic state task with explicit instructions to encode or retrieve, we have shown that younger adults can selectively engage in encoding and retrieval states in response to top-down instructions (Smith et al., 2022). Engagement in the retrieval state can impair later memory for to-be-encoded items (Long & Kuhl, 2019), impacts how stimuli are processed (Long & Kuhl, 2021), and coincides with a pattern of voltage topography consistent with DMN recruitment (Hong, Moore, Smith, & Long, 2023). Thus, younger adults recruit a DMN-like neural pattern when selectively engaging the retrieval state which has downstream consequences on behavior.

Mnemonic brain state engagement changes in healthy aging. We have found that older adults, like younger adults, can follow explicit instructions to engage in encoding or retrieval (Moore et al., 2025). Furthermore, mnemonic brain states can be reliably decoded in both age groups. Critically, however, the degree to which older adults engage mnemonic brain states differs from younger adults. In older adults, the neural activity patterns associated with encoding and retrieval are less robustly expressed and compared to younger adults, older adults display brief retrieval-like engagement when cued to encode. Our interpretation is that older adults may engage in retrieval regardless of its goal-relevance and require a minimum amount of time to switch out of this goal-irrelevant brain state.

Our hypothesis is that older adults are both biased towards retrieval and impaired at switching out of a task-irrelevant retrieval state. To test our hypothesis, we conducted a mnemonic state task in which we explicitly biased participants to engage mnemonic brain states while we recorded scalp electroencephalography (EEG). Critically, we manipulated the stimulus duration (the length of time that the stimulus was presented on screen) to determine how varying stimulus processing time would impact mnemonic brain state engagement for each age group. We used cross-study multivariate pattern analysis (MVPA) to assess mnemonic state evidence over time. We predicted that increased stimulus processing time (i.e. longer stimulus durations) would enable older adults to switch out of a task-irrelevant state and switch into the task-relevant mnemonic state.

## 2 Materials and Methods

### 2.1 Participants

Fifty younger adult (32 female, age range = 18-32, mean age = 20.34 years) and fifty older adult (38 female, age range = 60-87, mean age = 70.55 years) fluent English speakers from the University of Virginia (UVA) community participated. Younger adults (YAs) were recruited from flyers posted around the UVA campus and surrounding areas. Older adults (OAs) were recruited through the Virginia Cognitive Aging Project, a longitudinal study of adults (Salthouse, 2011). Our sample size was determined a priori based on pilot data described in the pre-registration report of this study (https://osf.io/n5jt3). All participants had normal or corrected-to-normal vision. Informed consent was obtained in accordance with the UVA Institutional Review Boards for Social and Behavioral Research and Health Sciences Research, and participants were compensated for their participation. All YA and OA participants completed the Montreal Cognitive Assessment (MoCA); any participant who scored less than 26 was excluded, based on previous literature (Nasreddine et al., 2005).

Six YA participants were excluded from the final dataset: five who scored lower than 26 on the MoCA and one additional participant whose recording was lost due to technical difficulties. Eight OA participants were excluded from the final dataset: one who had poor task performance across all conditions (memory accuracy *<* 2.5*SDs of the mean of the full OA dataset) and seven OAs who failed the MoCA. Thus, data are reported for the remaining 44 YA and 42 OA participants. The raw, de-identified data and the associated experimental and analysis codes used in this study will be made available via the Long Term Memory laboratory website (https://longtermmemorylab.com) upon publication.

### 2.2 Mnemonic State Task Experimental Design

Stimuli consisted of 648 object pictures, drawn from an image database with multiple exemplars per object category (Konkle, Brady, Alvarez, & Oliva, 2010), e.g. ‘apple’, ‘suitcase’, ‘bench’, ‘blender’). From this database, we randomly chose 162 unique object categories and four exemplars from each category. For each participant, one exemplar in a set of four served as a List 1 object, one as a List 2 object, and the two remaining exemplars served as lures for the recognition phase. Object condition assignment was randomly generated for each participant.

#### General Overview

The task design follows our previous work (Smith et al., 2022) with one critical manipulation. In each of nine runs, participants viewed two lists containing 18 object images. For the first list, each object was new (List 1 objects). For the second list (List 2 objects), each object was again new, but also categorically related to a List 1 object. For example, if List 1 contained an image of a suitcase, List 2 would contain an image of a different suitcase (Figure 1). Participants were explicitly directed to try to remember the images for a memory test prior to studying the List 1 and List 2 items. During List 1, participants were instructed to encode each new object. During List 2, however, each trial contained an instruction to either encode the current object (e.g., the new suitcase) or to retrieve the corresponding object from List 1 (the old suitcase). No behavioral responses were made in either List 1 or List 2. The critical manipulation was the duration of the object stimulus in List 2, which dictates the time that participants viewed the stimulus and had the opportunity to either encode the presented object or retrieve the corresponding List 1 object. Stimulus duration varied between 1000, 2000, and 4000 ms, and varied across runs, such that all List 2 trials within a run shared the same stimulus duration. Following nine runs, participants completed a two-alternative forced-choice recognition test that assessed memory for all List 1 and List 2 objects.

**Figure 1.**
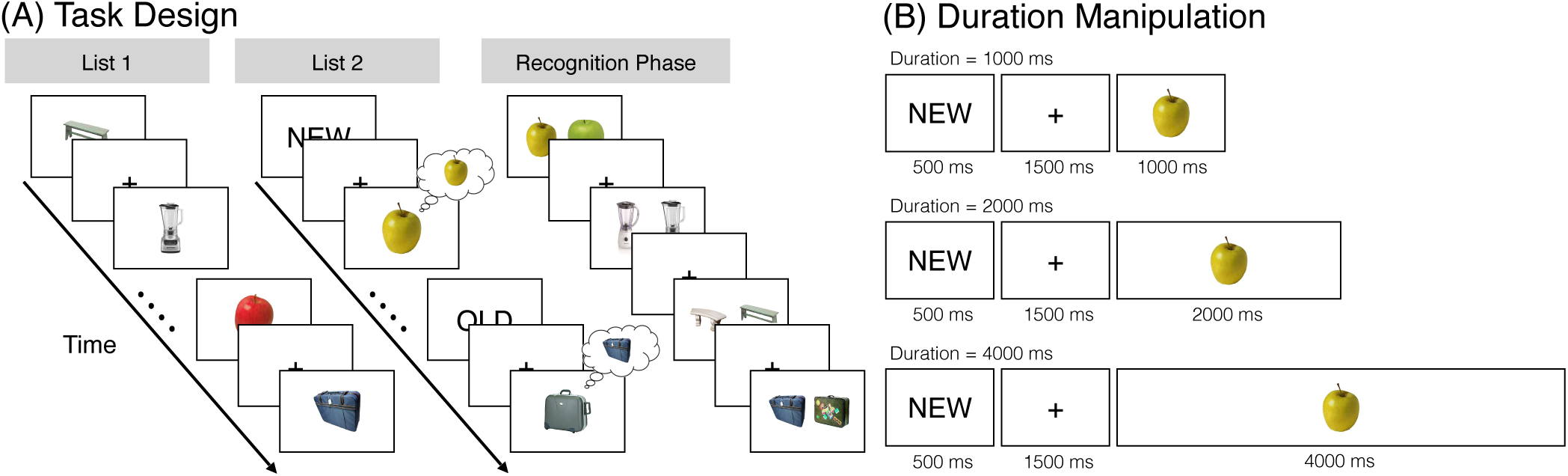
Task Design. **A** During List 1, participants studied individual common objects (e.g., blender, apple) that were each presented for 2000 ms. During List 2, participants saw novel objects that were categorically related to objects shown in List 1 (e.g., a new blender, a new apple). Preceding each List 2 object was an OLD instruction cue or NEW instruction cue. The OLD cue signaled that participants were to retrieve the corresponding object from List 1 (e.g., the old apple). The NEW cue signaled that participants were to encode the current object (e.g., the new apple). During List 2 trials, we manipulated the duration of the object stimulus and varied the stimulus duration conditions across runs, such that all List 2 trials within a run shared the same duration condition. Each run of the experiment contained a List 1 and List 2; object categories (e.g., blender) were not repeated across runs. Following nine runs, participants completed a two-alternative forced-choice recognition test that assessed memory for each List 1 and List 2 object. On each trial, a previously presented object, either from List 1 or List 2, was shown alongside a novel lure from the same category. The participant’s task was to choose the previously presented object. List 1 and List 2 objects were never presented together. **B** There were three stimulus duration conditions: 1000 ms, 2000 ms, and 4000 ms. The NEW or OLD instruction cue was presented for 500 ms and followed by a 1500 ms fixation.

#### List 1

On each trial, participants saw a single object presented for 2000 ms followed by a 1000 ms interstimulus interval (ISI). Participants were instructed to study the presented object in anticipation of a later memory test.

#### List 2

On each trial, participants saw a cue word, either “OLD” or “NEW” for 500 ms followed by a 1500 ms fixation. The cue was followed by presentation of an object for either 1000, 2000, or 4000 ms, which was followed by a 1000 ms ISI. All objects in List 2 were non-identical exemplars drawn from the same category as the objects presented in the immediately preceding List 1. That is, if a participant saw a suitcase and an apple during List 1, a different suitcase and a different apple would be presented during List 2. On trials with a “NEW” instruction (encode trials), participants were to encode the presented object. On trials with an “OLD” instruction (retrieve trials), participants tried to retrieve the categorically related object from the preceding List 1. Importantly, this design prevented participants from completely ignoring List 2 objects following “OLD” instructions in that they could only identify the to-be-retrieved object category by processing the List 2 object.

Participants completed nine runs with two lists in each run (List 1, List 2). Participants viewed 18 objects per list, yielding a total of 324 object stimuli from 162 unique object categories. Participants did not make a behavioral response during either List 1 or List 2. Following the nine runs, participants completed a two-alternative forced choice recognition test.

#### Recognition Phase

Following the nine runs, participants completed the recognition phase. On each trial, participants saw two exemplars from the same object category (e.g. two suitcases; Figure 1). One object had previously been encountered either during List 1 or List 2. The other object was a lure and had not been presented during the experiment. Because both test probes were from the same object category, participants could not rely on familiarity or gist-level information to make their response (Brainerd & Reyna, 2002). Trials were self-paced and participants selected (via button press) the previously presented object. Trials were separated by a 1000 ms ISI. There were a total of 324 recognition trials (corresponding to the 324 total List 1 and 2 objects presented in the experiment). Additionally, List 1 and List 2 objects never appeared in the same trial together, thus participants never had to choose between two previously presented objects. List 1 and List 2 objects were presented randomly throughout the test phase.

### 2.3 EEG Data Acquisition and Preprocessing

All acquisition and preprocessing methods are based on our previous work (Smith et al., 2022); for clarity we use the same text as previously reported. Electroencephalography (EEG) recordings were collected using a BrainVision system and an ActiCap equipped with 64 Ag/AgCl active electrodes positioned according to the extended 10-20 system. All electrodes were digitized at a sampling rate of 1000 Hz and were referenced to electrode FCz. Offline, electrodes were later converted to an average reference. Impedances of all electrodes were kept below 50 kΩ. Electrodes that demonstrated high impedance or poor contact with the scalp were excluded from the average reference calculations; however, all electrodes were included in all subsequent analysis steps following re-referencing. Bad electrodes were determined by voltage thresholding.

Custom Python codes were used to process the EEG data. We applied a high pass filter at 0.1 Hz, followed by a notch filter at 60 Hz and harmonics of 60 Hz to each participant’s raw EEG data. We then performed three preprocessing steps (Nolan, Whelan, & Reilly, 2010) to identify electrodes with severe artifacts. First, we calculated the mean correlation between each electrode and all other electrodes as electrodes should be moderately correlated with other electrodes due to volume conduction. We *z*-scored these means across electrodes and rejected electrodes with *z*-scores less than -3. Second, we calculated the variance for each electrode as electrodes with very high or low variance across a session are likely dominated by noise or have poor contact with the scalp. We then *z*-scored variance across electrodes and rejected electrodes with a *|z| >* = 3. Finally, we expect many electrical signals to be autocorrelated, but signals generated by the brain versus noise are likely to have different forms of autocorrelation. Therefore, we calculated the Hurst exponent, a measure of long-range autocorrelation, for each electrode and rejected electrodes with a *|z| >* = 3. Electrodes marked as bad by this procedure were excluded from the average re-reference. We then calculated the average voltage across all remaining electrodes at each time sample and re-referenced the data by subtracting the average voltage from the filtered EEG data. We used wavelet-enhanced independent component analysis (Castellanos & Makarov, 2006) to remove artifacts from eyeblinks and saccades.

### 2.4 EEG Data Analysis

We applied the Morlet wavelet transform (wave number 6) to the entire EEG time series across electrodes, for each of 46 logarithmically spaced frequencies (2-100 Hz; Smith et al., 2022). After log-transforming the power, we downsampled the data by taking a moving average across 100 ms time intervals from 4000 ms preceding to 4500 ms following object presentation during List 1 and List 2. We slid the window every 25 ms, resulting in 337 time intervals (85 non-overlapping). Power values were then *z*-transformed by subtracting the mean and dividing by the standard deviation power. Mean and standard deviation power were calculated across all List 1 and List 2 objects and across time points for each frequency.

### 2.5 Pattern Classification Analyses

Pattern classification analyses were performed using penalized (L2) logistic regression implemented via the sklearn linear model module in Python and custom Python code (Smith & Long, 2024). For all classification analyses, classifier features were composed of 63 electrodes and 46 frequencies. Before pattern classification analyses were performed, we completed an additional round of *z*-scoring across features (electrodes and frequencies) to eliminate trial-level differences in spectral power (Smith et al., 2022). Thus, mean univariate activity matched for all conditions and trial types.

#### 2.5.1 Within-Participant Mnemonic State Decoding

We performed leave-one-run-out cross-validated classification for each participant (penalty parameter = 1). We assessed classifier performance via “classification accuracy” and “classification evidence.” “Classification accuracy” represents a binary coding of whether the classifier successfully guesses the instruction condition. We used classification accuracy for general assessment of classifier performance (i.e., whether encode/retrieve instructions could be decoded). “Classification evidence” is a continuous value depicting the logit-transformed probability that the classifier assigns the correct mnemonic label (encode, retrieve) to each trial. Classification evidence was used as a trial-specific, continuous measure of mnemonic state information, which we used to determine the degree to which mnemonic state engagement varied across time as a function of the task instruction (encode, retrieve) and age group (younger, older). Due to the manner in which our classifier was configured, positive classification evidence values indicate more evidence for a retrieval state and negative classification evidence values indicate more evidence for an encoding state. This approach reflects our assumption – based on theoretical models of mnemonic brain states (Hasselmo et al., 2002; Hasselmo, 2005) – that encoding and retrieval exist along a continuum.

#### 2.5.2 Cross-Study Mnemonic State Decoding

We developed and validated a mnemonic state classifier to test mnemonic state engagement across YA and OA groups with a single training set, as within-participant classification can be driven by different features for different participants. We conducted three stages of classification using the same methods as in our prior work (Long, 2023; Smith & Long, 2024; Wheelock & Long, 2024; Han & Long, 2025). First, we conducted within participant leave-one-run-out cross-validated classification (penalty parameter = 1) on an independent set of younger adult participants who completed the mnemonic state task (N = 143, see ref. Hong et al., 2023; Han & Long, 2025 for details). The classifier was trained to distinguish encoding vs. retrieval states based on spectral power averaged across the 2000 ms stimulus interval during List 2 trials. We generated true and null classification accuracy values, as well as permuted condition labels (encode, retrieve) for 1000 iterations to generate a null distribution for each participant. Any participant whose true classification accuracy fell above the 90th percentile of their respective null distribution was selected for further analysis (N = 57). Second, we conducted leave-one-participant-out cross-validated classification (penalty parameter = 0.0001) on the selected participants to validate the mnemonic state classifier and obtained classification accuracy of 59.29% which is significantly above chance (*t* _56_ = 7.667, *p <* 0.0001), indicating that the cross-participant mnemonic state classifier is able to distinguish encoding and retrieval states. Finally, we applied the cross-participant mnemonic state classifier to the List 2 data in the current study, specifically in 100 ms intervals across the stimulus interval (1000, 2000, or 4000 ms). We extracted classification evidence, the logit-transformed probability that the classifier assigned a given trial a label of encoding or retrieval.

### 2.6 Statistical Analyses

We used mixed-effects ANOVAs and *t* -tests to assess the effect of age, instruction, and stimulus duration on memory accuracy and mnemonic state engagement. We used false discovery rate (FDR; Benjamini & Hochberg, 1995) to correct for multiple comparisons.

We used paired-sample *t* -tests to compare classification accuracy across participants to chance decoding accuracy, as determined by permutation procedures. Namely, for each participant, we shuffled the condition labels of interest (e.g., encode and retrieve for the within-participant List 2 instruction classifiers) and then calculated classification accuracy. We repeated this procedure 1000 times for each participant and then averaged the 1000 shuffled accuracy values for each participant. These mean values were used as participant-specific empirically derived measures of chance accuracy.

## 3 Results

### 3.1 Older and younger adults switch between encoding and retrieval in response to instructions

We first sought to replicate our prior finding that both younger adults (YAs) and older adults (OAs) can selectively engage mnemonic states in response to top-down instructions (Moore et al., 2025). Based on our assumption that retrieving the past comes at the expense of encoding the present, we expected to find a significant list by instruction interaction such that recognition accuracy is greater for List 2 encode compared to List 2 retrieve trials and greater for List 1 retrieve compared to List 1 encode trials. Following our pre-registration, we conducted a 2 *×* 2 *×* 2 mixed effects ANOVA with instruction type (encode, retrieve), list (1, 2) and age group (younger, older) as factors and recognition accuracy as the dependent variable (Figure 2, middle column). We conducted this ANOVA specifically for trials with the 2000 ms stimulus duration as this replicates our prior work (Moore et al., 2025). We report the full ANOVA results in Table 1 and highlight key findings here. We find a significant interaction between list and instruction type driven by greater recognition accuracy for List 2 encode trials (M = 0.8234, SD = 0.1106) compared to List 2 retrieve trials (M = 0.7808, SD = 0.1037; *t* _85_ = 3.275, *p* = 0.0015, *d* = 0.3978). The three-way interaction between age, instruction type, and list was not significant and Bayes Factor analysis revealed that a model without the three-way interaction term (H_0_) is preferred to a model with the three-way interaction term (H_1_; H_10_ = 0.3489, anecdotal evidence for H_0_). These results replicate our prior findings that both younger and older adults can selectively engage mnemonic states in response to explicit instructions.

**Figure 2.**
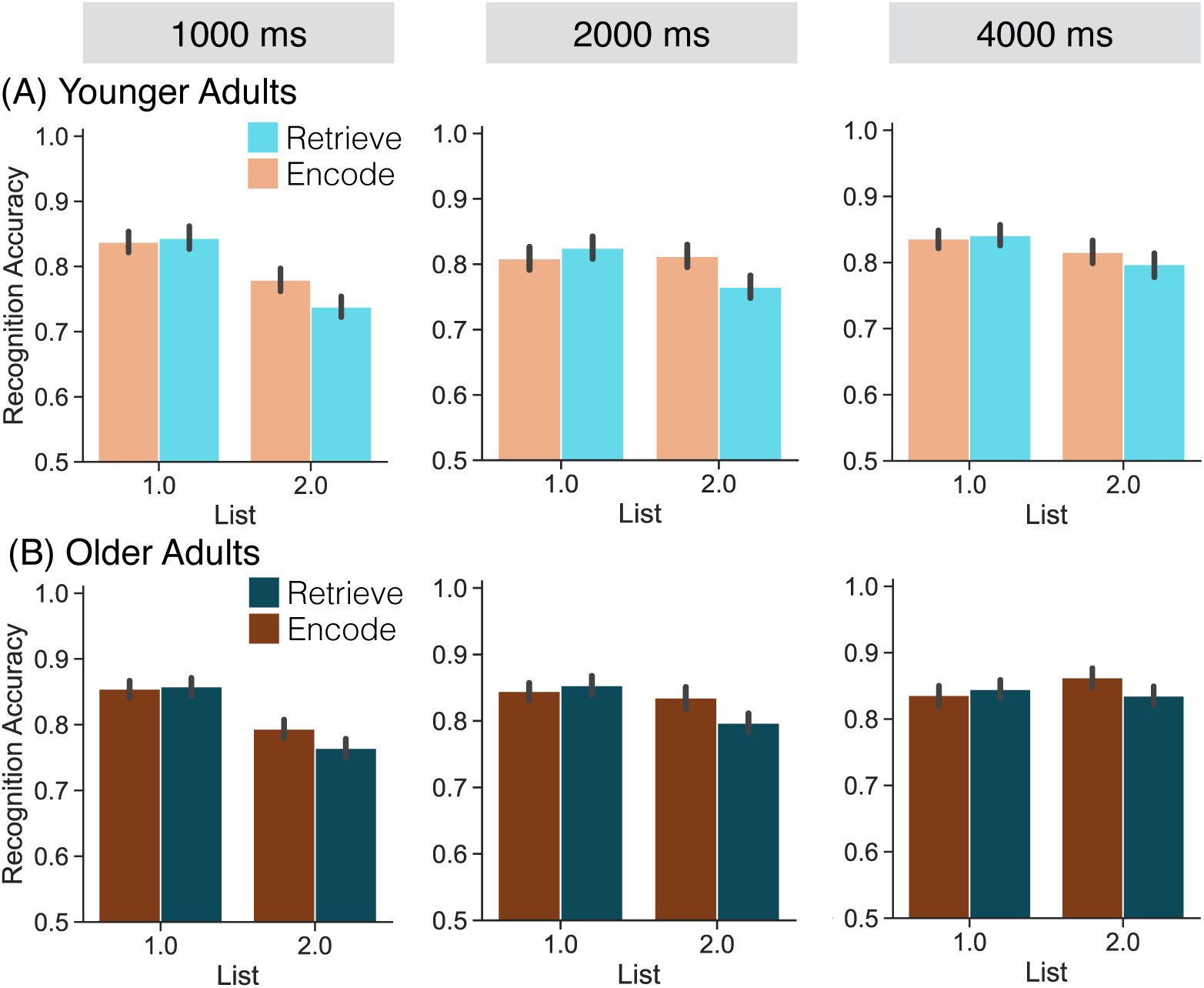
Recognition accuracy as a function of list, instruction, duration, and age. We assessed recognition accuracy as a function of list (1, 2), instruction (orange, encode; teal, retrieve), and duration (1000 ms, 2000 ms, 4000 ms) separately for each age group (younger adults, **A**; older adults, **B**). We find a significant interaction between list and instruction (*p* = 0.0015) with greater accuracy for List 2 objects paired with the encode, rather than retrieve, instruction. Error bars reflect standard error of the mean.

**Table 1.** Analysis of variance for the effect of age group, instruction type, and list on recognition accuracy for the 2000 ms duration condition.

| Effect | df | *F* | *p* | $\eta_p^2$ |
| --- | --- | --- | --- | --- |
| Main effect of age | 1,84 | 2.887 | 0.093 | 0.033 |
| **Main effect of instruction** | **1,84** | **4.019** | **0.0482** | **0.046** |
| **Main effect of list** | **1,84** | **12.56** | **0.0006** | **0.13** |
| Interaction of age × instruction | 1,84 | 0.005 | 0.9460 | < 0.0001 |
| Interaction of list × age | 1,84 | 0.08 | 0.7785 | 0.0009 |
| **Interaction of list × instruction** | **1,84** | **8.364** | **0.0049** | **0.091** |
| Interaction of age × list × instruction | 1,84 | 0.185 | 0.6683 | 0.002 |

Note: Bold values indicate $p < 0.05$

### 3.2 Longer stimulus duration facilitates subsequent memory

Having replicated our prior finding that both YAs and OAs can selectively encode and retrieve stimuli when given 2000 ms to do so, we next sought to assess how this selectivity would be impacted by shorter (1000 ms) vs. longer (4000 ms) stimulus durations (Figure 2, left and right columns). We expected that duration would influence older adults’ ability to switch out of a task-irrelevant retrieval state and into a task-relevant encoding state, meaning that the largest effects should be observed specifically for List 2 encode items. Based on this expectation, we specifically assessed memory accuracy for List 2 items and anticipated a significant three-way interaction between instruction, duration, and age, with older adults showing increased memory accuracy for List 2 encode trials as duration increases.

Following our pre-registration, we conducted a 2 *×* 2 *×* 3 ANOVA with instruction (encode, retrieve), age (younger, older), and stimulus duration (1000 ms, 2000 ms, 4000 ms) as factors and recognition accuracy as the dependent variable. We report the full ANOVA results in Table 2 and highlight key findings here. We find a main effect of stimulus duration whereby accuracy increases as a function of duration (1000 ms, M = 0.7687, SD = 0.0829; 2000 ms, M = 0.8021, SD = 0.0886; 4000 ms, M = 0.8271, SD = 0.0911). Although we find a main effect of instruction whereby memory accuracy is significantly greater for List 2 encode (M = 0.8161, SD = 0.0923) compared to List 2 retrieve (M = 0.7825, SD = 0.0787) trials, instruction type did not interact with age and/or duration. Bayes factor analysis revealed that a model without the three-way interaction term (H_0_) is preferred to a model with the three-way interaction term (H_1_; H_10_ = 0.0763, strong evidence for H_0_). Thus, we do not find evidence that stimulus duration modulates the ability to selectively engage mnemonic states. Instead, both younger and older adults are better at remembering List 2 encode vs. retrieve trials regardless of duration.

**Table 2.** Analysis of variance for the effect of age group, instruction type, and stimulus duration on recognition accuracy separately for List 2 and List 1 items.

| Effect | df | List 2 |  |  | List 1 |  |  |
| --- | --- | --- | --- | --- | --- | --- | --- |
| | | *F* | *p* | $\eta_p^2$ | *F* | *p* | $\eta_p^2$ |
| Main effect of age | 1,84 | 3.402 | 0.0686 | 0.0389 | 0.9606 | 0.3299 | 0.0113 |
| Main effect of instruction | 1,84 | **15.89** | **0.0001** | **0.1590** | 1.356 | 0.2475 | 0.0159 |
| Main effect of duration | 2,168 | **27.02** | **< 0.0001** | **0.2433** | 1.639 | 0.1972 | 0.0191 |
| Interaction of age × instruction | 1,84 | 0.0615 | 0.8047 | 0.0007 | 0.0194 | 0.8895 | 0.0002 |
| Interaction of duration × age | 2,168 | 1.002 | 0.3694 | 0.0118 | 1.557 | 0.2138 | 0.0182 |
| Interaction of duration × instruction | 2,168 | 0.6731 | 0.5115 | 0.0079 | 0.1757 | 0.8390 | 0.0021 |
| Interaction of age × duration × instruction | 2,168 | 0.2161 | 0.8059 | 0.0026 | 0.0825 | 0.9209 | 0.0010 |

Note: Bold values indicate $p < 0.05$

Our next goal was to assess the impact of instruction, age, and stimulus duration specifically on List 1 memory performance. Although we typically do not find strong List 1 modulations (e.g. Smith et al., 2022), we reasoned that we may find modulations when the stimulus duration changes – as shorter/longer durations would allow less/more time for List 1 retrieval. Following our pre-registration, we conducted a 2 *×* 2 *×* 3 ANOVA with instruction (encode, retrieve), age (younger, older), and stimulus duration (1000 ms, 2000 ms, 4000 ms) as factors and recognition accuracy as the dependent variable. We report the results of this ANOVA in Table 2; however, we did not find any significant effects or interactions. As in prior work, we find little impact of instruction on List 1 memory which in turn does not appear to be modulated by age or duration.

### 3.3 Older and younger adults engage mnemonic states in response to top-down instructions

We first sought to replicate our prior work (Moore et al., 2025) and assess the time course and strength of retrieval state engagement specifically for the 2000 ms stimulus duration. Based on our hypothesis that older adults are both biased toward retrieval and impaired at switching out of a task-irrelevant retrieval state, we expected to find a main effect of age whereby older adults show overall greater levels of retrieval evidence than younger adults. Based on our prior work (Moore et al., 2025), we expected to find a main effect of instruction and no significant interaction between age and instruction. Specifically, we expected to find that both younger and older adults can selectively engage mnemonic states in response to instruction such that both age groups would show more positive mnemonic state evidence (more evidence for the retrieval state) for retrieve compared to encode trials. To test our hypothesis, we utilized a cross-study mnemonic state classifier that was trained on previously collected data from younger adult participants (N=57) who completed the same mnemonic state task with a fixed stimulus duration of 2000 ms.

Following our pre-registration, we conducted a 2 *×* 2 *×* 20 mixed-effects ANOVA with instruction (encode, retrieve), age (younger, older), and time interval (twenty 100 ms intervals) as factors and mnemonic state evidence as the dependent variable (Figure 3, middle row). We report the results of this ANOVA in Table 3 and highlight the key findings here. We did not find a significant main effect of age. We find a significant main effect of instruction driven by greater mnemonic state evidence (i.e. evidence for the retrieval state) on retrieve (M = 0.0003, SD = 0.0648) compared to encode (M = -0.0271, SD = 0.0605) trials. The interaction between age and instruction was not significant; however, Bayes Factor analysis revealed that a model without the age*×*instruction interaction term (H_0_) is only slightly preferred to a model with the age*×*instruction interaction term (H_1_; H_10_ = 0.97, anecdotal evidence for H_0_). The lack of strong evidence either for or against an age by instruction interaction may be related to the marginal three-way interaction between age, instruction, and time interval, as well as our finding that younger adults show more late-interval (1000-2000 ms) mnemonic state evidence on retrieve trials (M = 0.0765, SD = 0.0858) relative to older adults (M = 0.0363, SD = 0.0914). Thus, if anything, older adults show less task-relevant (rather than irrelevant) retrieval state engagement relative to younger adults.

**Figure 3.**
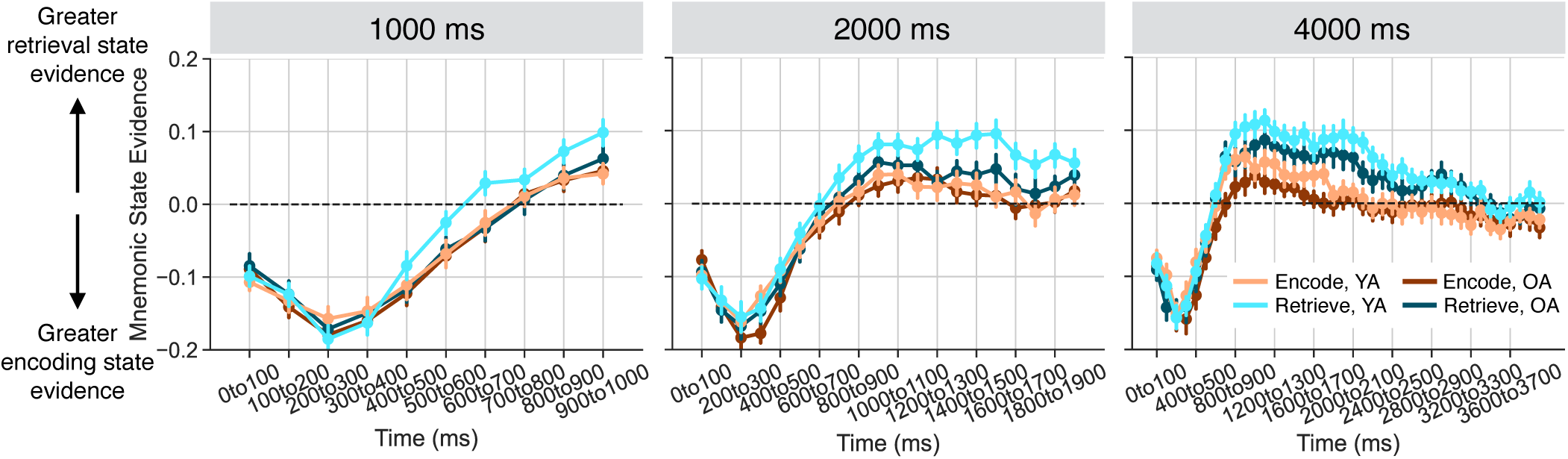
Mnemonic state evidence over time as a function of instruction, duration, and age. We applied a cross study classifier to the stimulus interval (twenty 100 ms time intervals) to measure mnemonic state evidence during List 2 as a function of instruction (encode, orange; retrieve, teal) and age (younger, YA; older, OA) separately for each stimulus duration condition (1000 ms, 2000 ms, 4000 ms). We find a significant main effect of instruction across all stimulus duration conditions (*p <* 0.0001) with greater mnemonic state evidence for retrieve compared to encode trials. Error bars reflect standard error of the mean.

**Table 3.** Mnemonic state evidence as a function of instruction and age for each duration condition.

| Effect | 1000 ms |  |  |  | 2000 ms |  |  |  | 4000 ms |  |  |  |
| --- | --- | --- | --- | --- | --- | --- | --- | --- | --- | --- | --- | --- |
| | df | *F* | *p* | $\eta_p^2$ | df | *F* | *p* | $\eta_p^2$ | df | *F* | *p* | $\eta_p^2$ |
| Main effect of age | 1,84 | 0.6385 | 0.4265 | 0.0075 | 1,84 | 2.720 | 0.1028 | 0.0314 | 1,84 | 1.998 | 0.1612 | 0.0232 |
| Main effect of time | **9,756** | **125.9** | **< 0.0001** | **0.5996** | **19,1596** | **97.52** | **< 0.0001** | **0.5373** | **39,3276** | **64.17** | **< 0.0001** | **0.4331** |
| Main effect of inst. | 1,84 | 2.917 | 0.0913 | 0.0336 | **1,84** | **11.75** | **0.0009** | **0.1227** | **1,84** | **33.90** | **< 0.0001** | **0.2875** |
| Int. of age × time | 9,756 | 0.8611 | 0.5599 | 0.0101 | 19,1596 | 0.5804 | 0.9222 | 0.0069 | 39,3276 | 0.8152 | 0.7859 | 0.0096 |
| Int. of time × inst. | **9,756** | **2.037** | **0.033** | **0.0237** | **19,1596** | **2.891** | **< 0.0001** | **0.0333** | **39,3276** | **3.318** | **< 0.0001** | **0.0380** |
| Int. of age × inst. | 1,84 | 0.7371 | 0.3930 | 0.0087 | 1,84 | 1.548 | 0.2169 | 0.0181 | 1,84 | 0.0461 | 0.8305 | 0.0005 |
| Int. of age × time × inst. | **9,756** | **2.144** | **0.0240** | **0.0249** | 19,1596 | 1.528 | 0.0673 | 0.0179 | 39,3276 | 0.5973 | 0.9778 | 0.0071 |

Note: Bold values indicate $p < 0.05$

### 3.4 Reduced task-relevant retrieval state engagement in older adults for shorter stimulus durations

Our primary goal was to test the hypothesis that older adults are both biased toward retrieval and impaired at switching out of a task-irrelevant retrieval state. We expected that decreasing stimulus processing time would limit older adults’ ability to switch out of the retrieval state. To the extent that younger, but not older, adults can switch into the task-relevant brain state within 1000 ms, we should find less retrieval state engagement for younger adults on encode trials later in the stimulus interval relative to older adults, who should show elevated retrieval state engagement across time and instruction. However, given our findings for the 2000 ms duration condition above, we may instead find more retrieval state engagement for younger compared to older adults on retrieve trials.

Following our pre-registration, we conducted a 2 *×* 2 *×* 10 mixed-effects ANOVA with instruction (encode, retrieve), age (younger, older), and time interval (ten 100 ms intervals) as factors specifically for the 1000 ms stimulus duration (Figure 3, top row). We report the results of this ANOVA in Table 3; critically, we find a significant three-way interaction between age, instruction and time.

We conducted two follow-up post-hoc mixed-effects ANOVAs with age (younger, older) and time interval (ten 100 ms intervals) as factors, separately for the encode and retrieve instruction trials. For the encode trials, we do not find a significant main effect of age (*F* _1,84_ = 0.0637, *p* = 0.8013, *η_p_*^2^= 0.0008). We find a significant main effect of time interval (*F* _9,756_ = 82.78, *p <* 0.0001, *η_p_*^2^= 0.4964). We do not find a significant age by time interaction (*F* _9,756_ = 0.4, *p* = 0.9353, *η_p_*^2^= 0.0047). For the retrieve trials, we do not find a significant main effect of age (*F* _1,84_ = 1.285, *p* = 0.2602, *η_p_*^2^= 0.151). We find a significant main effect of time interval (*F* _9,756_ = 89.64, *p <* 0.0001, *η_p_*^2^= 0.5163) and a significant age by time interaction (*F* _9,756_ = 1.991, *p* = 0.0377, *η_p_*^2^= 0.0232). These results run counter to our expectation that older adults are biased to retrieval. Rather than find elevated mnemonic state evidence on encode trials for older relative to younger adults – as would be predicted by the retrieval bias account – we find diminished mnemonic state evidence on retrieve trials for older relative to younger adults. Thus, rather than automatically engage the retrieval state when task-irrelevant, older adults show less retrieval state engagement when task-relevant. Given the short stimulus duration, these results suggest that older adults have difficulty switching *into*, rather than out of, the retrieval state in response to task instructions.

### 3.5 Comparable mnemonic state engagement across age for longer stimulus durations

Having shown that decreased stimulus processing time appears to diminish task-relevant retrieval state engagement in older adults, we next sought to test the impact of increased stimulus processing time on mnemonic state engagement. To the extent that increased stimulus processing time enables older adults to fully switch into task-relevant mnemonic states, we expected that for trials with the longest stimulus duration (4000 ms), both older and younger adults would show robust task-relevant mnemonic state engagement.

Following our pre-registration, we conducted a 2 *×* 2 *×* 40 mixed-effects ANOVA with instruction (encode, retrieve), age (younger, older), and time interval (forty 100 ms intervals) as factors specifically for the 4000 ms stimulus duration. We report the results of this ANOVA in Table 3 and highlight the key findings here. We find a main effect of instruction driven by greater retrieval state evidence on retrieve (M = 0.0215, SD = 0.052) compared to encode (M = -0.0133, SD = 0.051) trials. We find no effects of or interactions with age. Bayes Factor analysis revealed that a model without the three-way interaction term (H_0_) is preferred to a model with the three-way interaction term (H_1_; H_10_ = 8.02e-7, extreme evidence for H_0_). Thus, with longer processing time, both younger and older adults show selective mnemonic state engagement.

### 3.6 Longer stimulus durations promote greater mnemonic state separation

We directly compared mnemonic state engagement across duration conditions by averaging retrieval state evidence across the stimulus interval separately for each stimulus duration condition. Following our pre-registration, we conducted a 2 *×* 2 *×* 3 mixed-effects ANOVA with instruction (encode, retrieve), age (younger, older), and stimulus duration (1000ms, 2000ms, 4000ms) as factors. We do not find a significant main effect of age (*F* _1,84_ = 2.268, *p* = 0.1358, *η_p_*^2^= 0.0263). We find a significant main effect of stimulus duration (*F* _2,168_ = 63.06, *p <* 0.0001, *η_p_*^2^= 0.4288) driven by greater mnemonic state evidence – that is, more evidence for the retrieval state – as stimulus duration increases. We find a significant main effect of instruction (*F* _1,84_ = 18.27, *p <* 0.0001, *η_p_*^2^= 0.1786) driven by greater mnemonic state evidence for retrieve (M = -0.0106, SD = 0.0516) compared to encode (M = -0.0361, SD = 0.0518) trials, consistent with the effects reported above. We do not find a significant interaction between either age and duration (*F* _2,168_ = 0.1590, *p* = 0.8531, *η_p_*^2^= 0.0019) or age and instruction (*F* _1,84_ = 1.060, *p* = 0.3063, *η_p_*^2^= 0.0125). We find a significant interaction between instruction and duration (*F* _2,168_ = 3.431, *p* = 0.0346, *η_p_*^2^= 0.0392). The three-way interaction between age, instruction, and duration was not significant (*F* _2,168_ = 0.6247, *p* = 0.5366, *η_p_*^2^= 0.0074). Bayes Factor analysis revealed that a model without the three-way interaction term (H_0_) is preferred to a model with the three-way interaction term (H_1_; H_10_ = 0.09, strong evidence for H_0_). We conducted three follow-up post-hoc paired *t* -tests to investigate the instruction by duration interaction (Figure 4). First, we subtracted mnemonic state evidence on encode trials from mnemonic state evidence on retrieve trials to obtain a difference score whereby positive values indicate greater mnemonic state evidence (i.e. evidence for the retrieval state) for retrieve relative to encode trials. Next, we performed pairwise comparisons of these difference scores across durations. Difference scores were significantly different between the 1000 ms (M = 0.0143, SD = 0.0769) and 4000 ms (M = 0.0348, SD = 0.0547) conditions, whereby the dissociation between retrieve and encode instructions was significantly greater for the longer duration condition (*t* _85_ = 2.620, *p* = 0.0104, *d* = 0.307, FDR corrected). Difference scores did not significantly differ between 1000 ms and 2000 ms (M = 0.0274, SD = 0.074; *t* _85_ = 1.496, *p* = 0.1383, *d* = 0.1744) or between 2000 ms and 4000 ms (*t* _85_ = 1.045, *p* = 0.2988, *d* = 0.1126).

**Figure 4.**
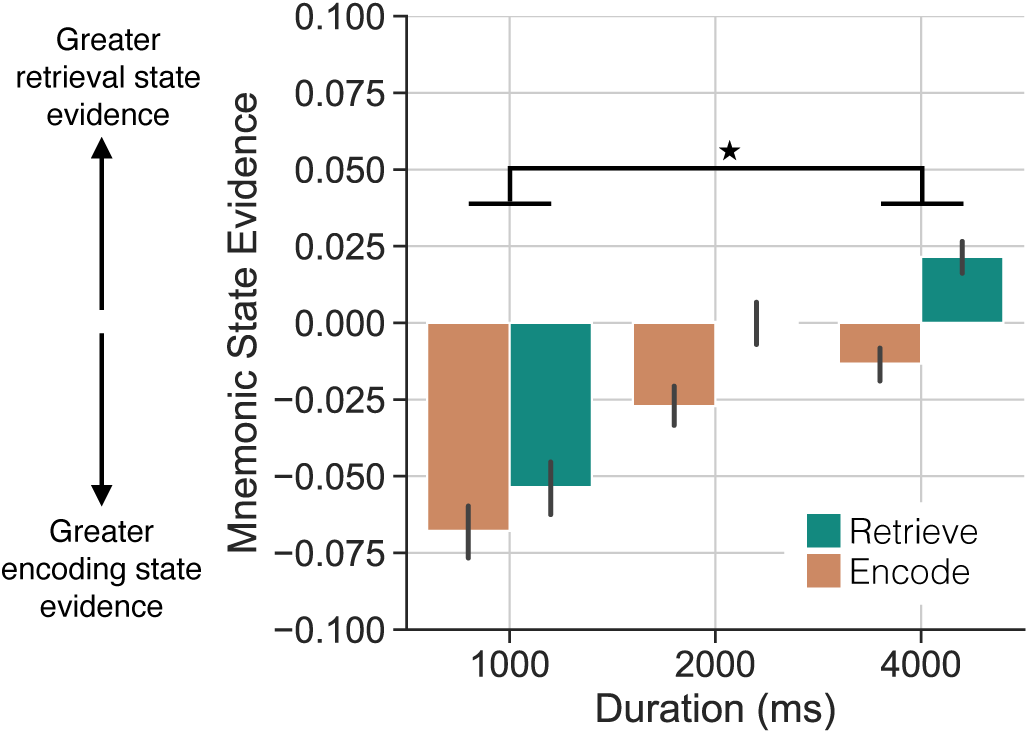
Mnemonic state evidence as a function of instruction and duration. We averaged mnemonic state evidence across time interval and age and compared encode (orange) to retrieve (teal) instructions across each stimulus duration condition. The dissociation between encode and retrieve trials was significantly greater for the longest duration (4000 ms) compared to the shortest duration (1000 ms). Error bars reflect standard error of the mean. \**p <* 0.05, FDR corrected.

Together, our results suggest that both younger and older adults are better able to engage task-relevant mnemonic brain states as the time to do so increases, and that older adults are selectively impaired at engaging in retrieval under short stimulus durations.

### 3.7 Evidence for an age-related retrieval state bias

In the preceding planned analyses, we performed cross-study decoding with an independent dataset to assess the impact of stimulus duration on mnemonic state engagement across age. We chose this approach to avoid potential confounds driven by stimulus duration and to equate classifier training across younger and older adult participants. Within-participant decoding requires separate classifiers for each duration condition, which would reduce the overall number of trials available for training (n = 36 total trials, 18 per class). Failure to find above chance decoding in this case could be attributed to low trial counts, rather than a true lack of difference in the neural patterns during encode vs. retrieve trials. Additionally, although classification accuracy may exceed chance in both younger and older adults, the features used to generate these values may differ across age groups. Using the same training dataset avoids both of these issues. However, to the extent that mnemonic states differ across individuals – and especially, across age groups – we anticipated that it may be necessary to conduct within-participant decoding. Given the lack of an impact of stimulus duration on behavior and the relatively modest effect of stimulus duration on mnemonic state engagement – and not in the predicted direction – we next performed a series of within-participant leave-one-run-out cross-validated decoding analyses.

We first assessed classification accuracy, or how well encode vs. retrieve instructions can be distinguished, separately for each age group and stimulus duration (Figure 5). We expected to find above chance classification across all stimulus durations for younger adults as they should be able to engage task-relevant mnemonic states even when the stimulus duration is short (1000 ms). In contrast, we did not expect to find above chance classification in older adults specifically for the 1000 ms duration. For younger adults, we find significantly above chance classification accuracy (as determined by permutation procedure, see Methods) for the 2000 ms and 4000 ms duration conditions, but not the 1000 ms duration condition (Table 4). For older adults, we find significantly above chance classification accuracy only for the 4000 ms duration condition.

**Figure 5.**
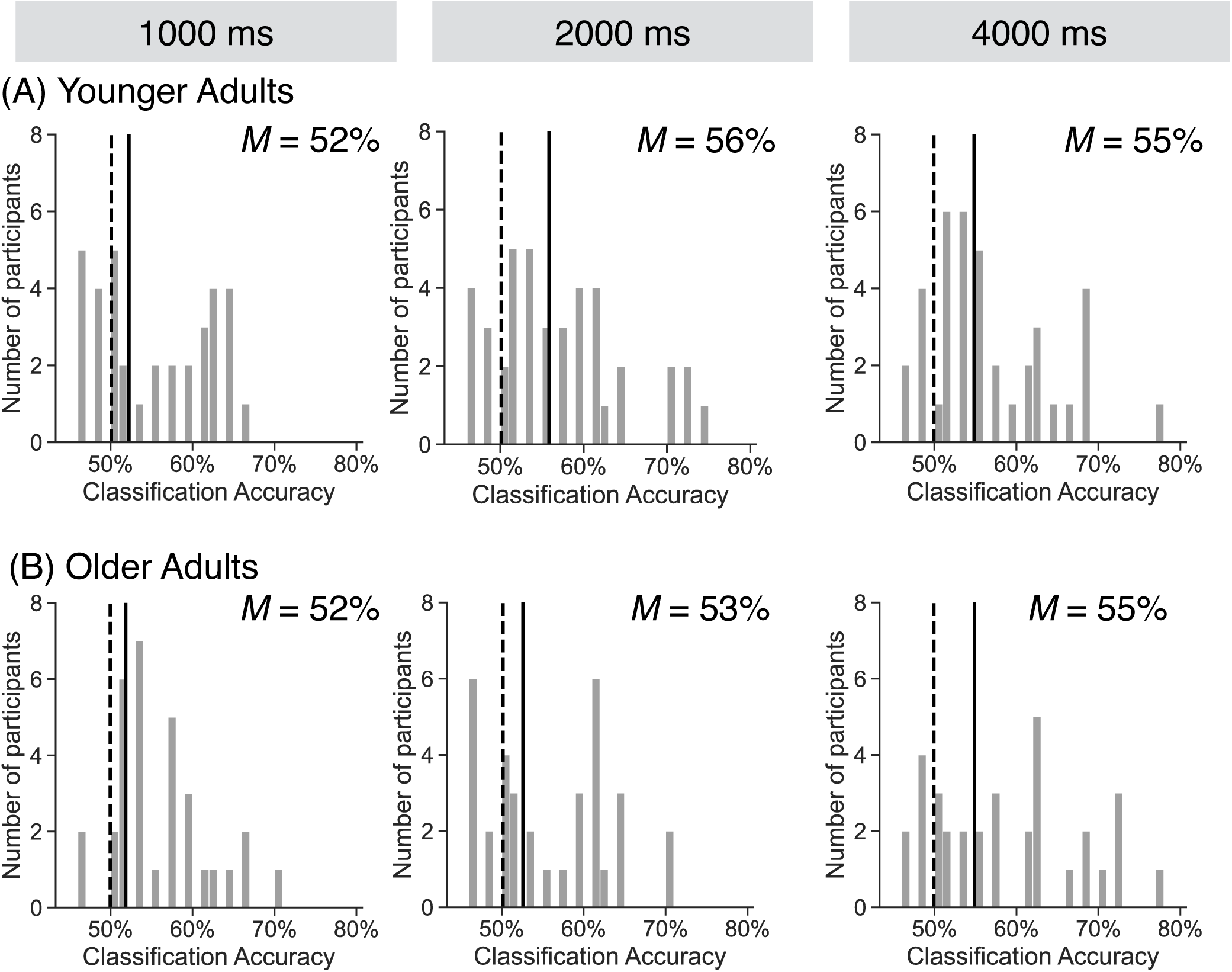
Classification accuracy as a function of duration and age group. We conducted within-participant leave-one-run-out cross-validated classification separately for each age group (younger adults, **A**; older adults, **B**) and duration (1000 ms, 2000 ms, 4000 ms). Histograms show distributions of classification accuracy values across participants with mean accuracy shown via the solid black vertical line. Chance is determined via permutation procedure and mean chance accuracy is shown via the dashed black vertical line. For the 1000 ms duration condition, mean classification accuracy did not differ significantly from chance for either age group (younger adults, *p* = 0.11; older adults, *p* = 0.19). For the 2000 ms duration condition, mean classification accuracy was significantly greater than chance for younger adults (*p <* 0.0001), but not for older adults (*p* = 0.0986). For the 4000 ms duration condition, mean classification accuracy was significantly greater than chance for both age groups (younger adults, *p* = 0.0007; older adults, *p* = 0.005).

**Table 4.** Within-participant mnemonic state decoding accuracy by age and duration.

|  |  | 1000 ms |  |  |  |  |  | 2000 ms |  |  |  |  |  | 4000 ms |  |  |  |  |  |
| --- | --- | --- | --- | --- | --- | --- | --- | --- | --- | --- | --- | --- | --- | --- | --- | --- | --- | --- | --- |
| Age group | df | M | SD | *t* | *p* | *d* | *n* | M | SD | *t* | *p* | *d* | *n* | M | SD | *t* | *p* | *d* | *n* |
| Younger | 43 | 52.23% | 8.74% | 1.621 | 0.11 | 0.3464 | 13 | **55.86%** | **8.16%** | **4.702** | **<0.0001** | **0.9933** | 13 | **54.84%** | **8.9%** | **3.648** | **0.0007** | **0.7831** | 12 |
| Older | 41 | 51.85% | 9.14% | 1.333 | 0.19 | 0.2925 | 7 | 52.6% | 9.23% | 1.690 | 0.0986 | 0.3728 | 12 | **54.89%** | **10.66%** | **2.967** | **0.005** | **0.6588** | 15 |

Note: Bold values indicate $p < 0.05$ . *n* indicates the number of participants with above chance accuracy values

Thus, the within-participant decoding analyses broadly mirror the results obtained from the application of the independent classifier in the planned analyses above. Mnemonic decoding in older adults does not exceed chance for shorter stimulus durations, presumably because these conditions provide insufficient time for older adults to shift into the instructed states. Critically, however, these classification accuracy results do not indicate the degree to which younger and older adults engage mnemonic states on retrieve vs. encode trials. A classifier may correctly label trials “encode” or “retrieve” for both younger and older adults; however, the confidence with which those labels are applied may differ across age. Specifically, whereas a classifier could assign the correct label (e.g. “retrieve”), the evidence for that label may be strong (approaching 1, prior to logit-transforming) or weak (approaching 0.5, prior to logit-transforming). Thus, in addition to assessing classification accuracy, we assessed classification evidence which can be interpreted as the classifier’s confidence in the label that it assigns to each trial. Greater distinguishability between spectral patterns for encode vs. retrieve trials should yield higher evidence values.

We measured mnemonic state evidence over time separately for each age group, instruction, and duration condition (Figure 6). We conducted three mixed-effects ANOVAs, one for each duration, with factors of instruction (encode, retrieve), age (younger, older), and time interval (ten, twenty, or forty 100 ms windows). We report the results of these ANOVAs in Table 5 and highlight the key findings here. Across all duration conditions, we find significant main effects of instruction, which is to be expected given that some participants across both age groups showed above chance within-participant mnemonic state decoding. For the 2000 ms duration condition, we find a significant interaction between age and time interval, indicating that the temporal dynamics of mnemonic state engagement differs across younger and older adults.

**Figure 6.**
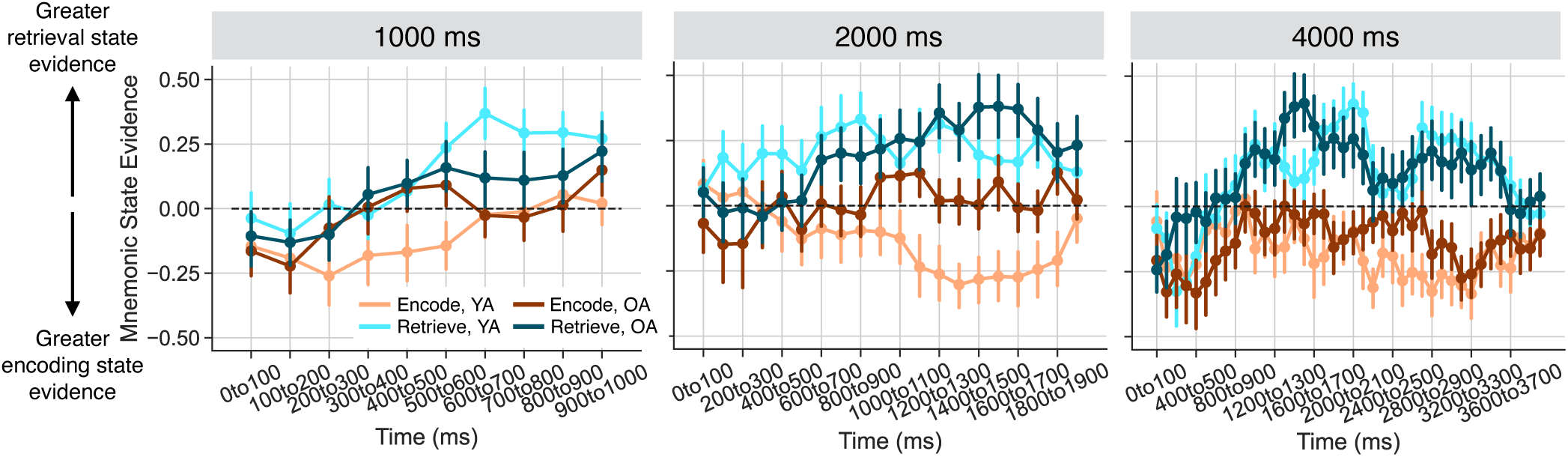
Within-participant derived mnemonic state evidence as a function of instruction, duration, and age. We conducted leave-one-run-out cross-validated classification separately for each age group (younger adults, YA; older adults, OA) and duration condition (1000 ms, 2000 ms, 4000 ms). Each within-participant classifier was trained on z-scored power averaged across the respective stimulus interval and tested on ten, twenty, or forty 100 ms time intervals. Mnemonic state evidence is shown as a function of age (YA, lighter hues; OA, darker hues), instruction (encode, orange; retrieve, teal), and time. Error bars reflect standard error of the mean.

**Table 5.**
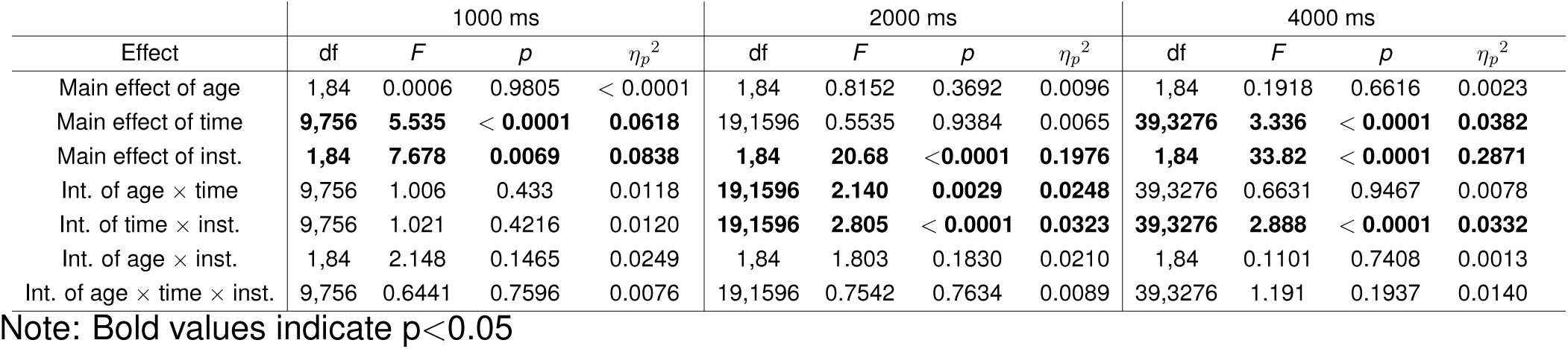
Within-participant-derived mnemonic state evidence as a function of instruction and age for each duration condition.

We conducted two follow-up post-hoc two-sample *t* -tests comparing younger and older adult mnemonic state engagement across early (0-1000ms) and late (1000-2000ms) in the stimulus interval for the 2000 ms duration condition. Mnemonic state engagement does not differ significantly between younger (M = 0.0811, SD = 0.41) and older (M = 0.0202, SD = 0.367) adults in the early stimulus interval (*t* _84_ = 0.7162, *p* = 0.4759, *d* = 0.1565). Mnemonic state engagement is significantly greater for older (M = 0.1752, SD = 0.4132) compared to younger (M = -0.009, SD = 0.3706) adults later in the stimulus interval (*t* _84_ = 2.153, *p* = 0.0342, *d* = 0.4694) meaning that regardless of instruction, older adults are more likely to engage the retrieval state than younger adults. Later in the stimulus interval, older adults show a significant increase in mnemonic state evidence relative to 0 (one-sample t-test vs. 0: *t* _41_ = 2.715, *p* = 0.0096, *d* = 0.424) whereas younger adults do not (one-sample t-test vs. 0: *t* _43_ = -0.1591, *p* = 0.8744, *d* = 0.0243). Thus, within-participant classification suggests an age-related retrieval state bias.

## 4 Discussion

The goal of the current study was to determine the extent to which stimulus processing time impacts older adult mnemonic state engagement. To test the hypothesis that older adults are both biased toward retrieval and impaired at switching out of a task-irrelevant retrieval state, we recorded scalp electroencephalography (EEG) while younger and older adult participants explicitly encoded and retrieved object stimuli under variable stimulus durations. We used an independently-validated mnemonic state classifier to measure mnemonic state engagement. Surprisingly, we find that compared to younger adults, older adults show a reduction in task-relevant retrieval state engagement on trials with the shortest stimulus duration. However, within-participant decoding revealed an age-related bias toward the retrieval state. Taken together, these findings suggest that although older adults retain the ability to engage encoding and retrieval states, they require more processing time to both initiate and maintain task-relevant mnemonic states.

Replicating prior behavioral work (Moore et al., 2025), we find that both younger and older adults can selectively switch between encoding and retrieval in response to top-down instructions. Specifically, we find that recognition accuracy is greater for items paired with the encode instruction relative to items paired with the retrieve instruction for both age groups. To the extent that older adults require more time to switch out of a task-irrelevant retrieval state and into a task-relevant encoding state, we expected memory performance for items paired with the encode instruction to increase as stimulus duration increased. However, we do not find evidence that stimulus duration significantly modulates the impact of instruction on memory performance in either age group. This pattern differs from the substantial literature on the processing speed hypothesis which posits that as age increases, the rate of cognitive processing decreases (Salthouse, 1996, 1993). Although we cannot draw strong conclusions from null results, it may be that the current task structure sufficiently supported older adults’ performance reducing the potential impact of processing time on behavior. Older adults’ episodic deficits are strongest when retrieval is self-initiated and unsupported relative to tasks that provide structure or support (Wingfield & Kahana, 2002; Logan, Sanders, Snyder, Morris, & Buckner, 2002). In the current study, the two-alternative forced choice recognition test provides participants with structured retrieval cues, reducing reliance on self-initiated search and strategic organization. Environmental supports during study and/or test can aid older adults’ ability to follow top-down instructions and can lead older adults to perform comparably to younger adults on recognition tests (Craik, 2022).

Despite comparable behavioral performance, we find that for the shortest duration condition, older adults display diminished retrieval state engagement on retrieve trials compared to younger adults. Based on evidence that older adults show a bias toward semantic memory (Wynn et al., 2020), have difficulty inhibiting retrieval (Healey, Ngo, & Hasher, 2014), and prior investigation of mnemonic states in aging (Moore et al., 2025), we expected older adults to automatically engage the retrieval state regardless of instruction. Furthermore, we specifically expected that a short, 1000 ms stimulus duration would provide insufficient time for older adults to switch out of the task-irrelevant retrieval state before encode trials ended. However, we find that for trials with a 1000 ms duration, older adults show less task-relevant retrieval state engagement relative to younger adults. This finding suggests that older adults may have difficulty switching *into*, rather than or in addition to out of, the retrieval state under time constraints. Indeed, older adults also show reduced task-relevant retrieval state engagement even when given more time (2000 ms). As age-related slowing in strategic encoding and retrieval processes puts older adults at a disadvantage when given the same time as younger adults (Foster & Giovanello, 2020; Chwiesko et al., 2023), longer processing time may help mitigate these disadvantages. Consistent with this account, with extended time (4000 ms) we find a robust dissociation between encode and retrieve trials and no age differences in mnemonic state engagement. When we directly compare the longest vs. shortest stimulus duration conditions, we find significantly greater dissociation between encode and retrieve trials for the longest duration across both age groups. Thus, with sufficient processing time, older adults are able to engage young-adult like mnemonic brain states to a similar degree as younger adults.

Our finding that older adults under-recruit the retrieval state in the context of time pressure runs counter to our hypothesis and prior findings. This result suggests that variability in stimulus timing may have impacted mnemonic state engagement. Importantly, the younger adult data also differ from past work. In prior studies utilizing a fixed stimulus duration, we found robust cross-study encoding state engagement on encode trials in younger adults (Moore et al., 2025; Han & Long, 2025). Here, however, we find that younger adults recruit the (task-irrelevant) retrieval state on encode trials. We used nearly the same mnemonic state task, recruited from the same participant database, and applied identical data analytic methods across both the current and prior studies. The central difference between the studies is variable (current) vs. fixed (prior) stimulus duration suggesting that uncertainty around stimulus timing may have influenced our results. Uncertainty can impact how representations are updated and how cognitive control is deployed (Mushtaq, Bland, & Schaefer, 2011) and older and younger adults respond differently to uncertainty (Kosciessa, Mayr, Lindenberger, & Garrett, 2024). Taken together, variability in stimulus duration, and possibly uncertainty more generally, may amplify the ‘default’ tendency to automatically engage the retrieval state – for both younger and older adults – due to a reduction in control resources. By design, all stimuli could be used as retrieval cues due to their categorical association to a previously presented stimulus. Contextual overlap between stimuli can result in younger adults automatically engaging the retrieval state (Smith et al., 2022). Thus, to the extent that some degree of control is deployed in response to uncertainty, there will be a reduction in control resources available to inhibit automatic retrieval, yielding the overall increase in retrieval state evidence across both younger and older adults observed in the current study. At the same time, the change in control demands due to uncertainty may also impinge on older adults’ ability to selectively engage task-relevant mnemonic states. These findings suggest that mnemonic states likely reflect a combination of automatic and controlled processes; identifying the unique contributions of each type of process is an important direction for future work.

We find above chance, but relatively modest, within-participant mnemonic state decoding which is also consistent with the idea that variable stimulus duration impacts mnemonic state engagement. To measure how stimulus duration impacts retrieval state engagement, we initially utilized an independently validated mnemonic state classifier. This classifier was trained on a separate group of younger adult participants who performed the mnemonic state task with a fixed stimulus duration (2000 ms). We used this approach rather than within participant decoding for two key reasons. First, the number of trials available for decoding in the current study is only 36 (18 per class) within each stimulus duration condition. Thus, failure to find above chance decoding may be due to low trial counts rather than true lack of dissociation between conditions. Second, utilizing the same training set for both younger and older adults ensures that age-related differences are not the result of differences in the information to which the classifier has access during training. However, the use of a classifier trained exclusively on younger adult data assumes that older adults will use the same mechanisms to perform the mnemonic state task which may or may not be the case. Due to reported differences between younger and older adult electrophysiology (e.g. Voytek et al., 2015), we additionally performed within-participant mnemonic state decoding. We find above chance within-participant mnemonic state decoding for both the 2000 ms and 4000 ms stimulus duration for younger adults, but only for the 4000 ms duration for older adults. Furthermore, the percentage of participants who show above chance decoding is relatively modest, especially compared with our prior work. This reduced decoding performance may be related to both the number of trials available for training as well as the potential duration-driven uncertainty effects discussed above.

Within-participant decoding revealed an age-related bias toward retrieval. Although classification accuracy may be significantly above chance for both younger and older adults, evidence in support of the classifier’s response can vary such that a classifier can be accurate and confident (large evidence values) or accurate but not confident (small evidence values). Evidence is expected to scale with the underlying fidelity of the neural patterns on which the classifier is tested with larger evidence values for higher fidelity patterns. We find that older adults show significantly elevated retrieval state evidence in the second half of the 2000 ms stimulus interval regardless of instruction whereas younger adults show no such elevation. Thus, together with the cross-study decoding results, we find that older adults under-recruit a young-adult-like task-relevant retrieval state and over-recruit the retrieval state when task-irrelevant. These brain state recruitment effects appear most prominently under time pressure.

Our interpretation is that age-related alterations in mnemonic state engagement are driven by the ability to initiate, maintain, and switch between task-relevant and task-irrelevant brain states as we found no age-related differences in any metric for the longest stimulus duration. This interpretation is consistent with the default-executive coupling hypothesis of aging (DECHA, Turner & Spreng, 2015; Spreng & Turner, 2019), which suggests that age-related cognitive changes arise from reduced prefrontal control signaling coupled with increased activity in the default mode network (DMN). We expect that diminished prefrontal control signals impact the ability to switch between mnemonic states. To the extent that exogenous, external attention is captured by presentation of the visual stimulus (Chun, Golomb, & Turk-Browne, 2011), older adults may have more difficulty disengaging this external attention to switch into an internally directed retrieval state. At the same time, due to hyperactivity within the DMN, a region known to support semantic processing (Binder et al., 2009) and internal mentation (Buckner & DiNicola, 2019), older adults may over-recruit the retrieval state when task-irrelevant, consistent with existing work showing an over-reliance on gist-level or semantic knowledge in healthy aging (Brod & Shing, 2019; Wynn et al., 2020; Matijevic et al., 2024). Counteracting such a bias also requires control, specifically inhibitory control, which tends to decrease with age (Amer et al., 2016; Sambataro et al., 2010). Thus, changes in executive functioning may account for changes in mnemonic state engagement. An important question for future work is the extent to mnemonic brain states are malleable – can experience with mnemonic brain state engagement facilitate both detecting task-irrelevant brain states and selecting task-relevant brain states?

Age-related changes in mnemonic state engagement likely impact how stimuli are represented. Prior work has suggested that older adults require more time to encode specific, verbatim representations similar to representations formed by younger adults (Greene & Naveh-Benjamin, 2024; Cowan et al., 2024). As mnemonic brain state engagement influences how stimuli are represented in younger adults (Long & Kuhl, 2021), over-recruitment of a task-irrelevant retrieval state may impair older adults’ ability to encode verbatim details. Similarly, altered mnemonic brain state engagement relate to the finding that neural representations become de-differentiated in healthy aging (Hill, King, & Rugg, 2021). Less time spent in task-relevant brain states may result in reduced distinctiveness of the to-be-encoded/retrieved stimuli (Koen & Rugg, 2019; Folville et al., 2020; Pauley, Karlsson, & Sander, 2024; Srokova, Aktas, Koen, & Rugg, 2024).

Taken together, these findings suggest that under time constraints, older adults both under-recruit task-relevant brain states and over-recruit task-irrelevant brain states. Thus, aging-related differences in memory performance may be the result of changes in the ability to initiate, maintain, and switch between task-relevant brain states. To the extent that mnemonic state engagement impacts downstream processing, age-related differences in executive processes that support mnemonic brain state engagement may account for memory deficits in healthy aging.

## 5 Acknowledgments

This work was supported by a grant from the National Institutes of Health (NINDS R01 NS132872, PI: N.M.L.) and a grant from the UVA Brain Institute (PI: N.M.L.).

## 6 Data and code availability

The datasets generated in the current study and all experimental codes used for data collection and data analysis will be made available on the Open Science Foundation (OSF) via the Long Term Memory Lab website (http://longtermmemorylab.com/publications/) upon publication.

## 7 CRediT authorship contribution statement

**Hannah R. Buras:** Data Curation, Formal analysis, Investigation, Software, Validation, Visualization, Writing – original draft, Writing – review & editing.

**Subin Han:** Data Curation, Formal analysis, Investigation, Writing – review & editing.

**Nicole M. Long:** Conceptualization, Formal analysis, Funding acquisition, Methodology, Project administration, Resources, Software, Supervision, Validation, Visualization, Writing – review and editing.

